# Establishment of *Galleria mellonella* as a Model for *Achromobacter xylosoxidans* Infection

**DOI:** 10.64898/2026.01.27.702115

**Authors:** Cameron Lloyd, Breanna Wimbush, Ngoc Thien Lam, Parker Adams, Pooja Acharya, Hanh Ngoc Lam

**Author notes:** **Correspondence:** Hanh Ngoc Lam.

## Abstract

*Achromobacter xylosoxidans (Ax)* is an emerging pathogen with a strong capacity to adapt to different niches, but its pathogenesis is poorly understood. To investigate the virulence of this versatile bacterium, alternative infection models are valuable. *Galleria mellonella* wax moth presents cost and ethical advantages as an *in vivo* infection model. Here, we investigate the utility of *Galleria* as a model of *Ax* infection and demonstrate that mortality following *Ax* infection in *Galleria* recapitulates survival outcomes observed in infected mice. We further show that the *Galleria* infection model can be used to examine antimicrobial activity against *Ax*. Visualization of hemocytes suggested that *Ax* was internalized into immune cells, similar to what is observed in vertebrate models. Overall, our work establishes *Galleria mellonella* as a model of *Ax* infection that mirrors disease severity and innate immune cell interactions in murine models.

## Introduction

*Achromobacter xylosoxidans* (*Ax*) is a gram-negative emerging opportunistic pathogen that is highly associated with immunocompromised patients (1-3). In addition to lung infections, *Ax* can cause skin and wound infections, urinary tract infections (UTIs), and systemic infections (4, 5). These infections can cause severe organ damage as a result of bacteremia and dissemination to immune-privileged sites, such as the central nervous system (6, 7). *Ax* can cause recurrent and prolonged infections and is highly resistant to antibiotic treatment (8-10). However, *Ax* has been largely understudied (11, 12).

To investigate *Ax* infection, *in vitro* cell based assays and *in vivo* mouse assays were used to model infection (11, 13). While *in vitro* models are useful for screening and characterizing virulence factors, they can fail to capture the complexity of *in vivo* infection. Indeed, infections are known to elicit different effects *in vivo* than those observed *in vitro* (14, 15). Thus, identification of an alternative model that recapitulates the *in vivo* murine model while reducing cost and ethical concerns would be beneficial.

*Galleria mellonella* is an invertebrate species that has gained increasing attention as an alternative to mammalian models of infection (16-19). Although *Galleria* lacks an adaptive immune response, its innate immune response is strikingly similar to that of mammals (20, 21). Therefore, *Galleria* has been employed to assess infection-associated mortality and treatment efficacy for related pathogens, including *Pseudomonas aeruginosa* and *Bordetella bronchiseptica* (22-25). *Galleria* has also become a widely used model for studying infection caused by a wide range of agents, from bacteria to fungi (26, 27). Development of a *Galleria mellonella* infection model would provide a cost-effective, time-efficient, and ethically improved alternative to murine models. Here, we describe the application of *Galleria mellonella* as an *in vivo* infection model by comparing infection outcomes in *Galleria* to those observed in a murine intratracheal instillation model and by demonstrating the use of this model in examining antibiotic treatment of *Ax* infection. We establish *Galleria* as an intermediary infection model for *Ax* that bridges *in vitro* assays and murine models.

## Methods

### Bacterial culture

*Achromobacter xylosoxidans* isolates (GN050, GN008, NIH-018 series) were obtained from previous work (28, 29). All *Ax* bacterial strains used in this study were first streaked onto LB Agar (LBA) containing no antibiotics and grown at 37°C overnight. Strains were then incubated at room temperature for 24 hours. Following incubation, bacteria were collected from the plates, suspended in phosphate-buffered saline (PBS) and optical density at 600nm (OD600) was measured using a spectrophotometer. The resulting suspension was diluted in PBS to the desired OD600 to obtain approximately 1x10^9^ CFU/mL and was serially diluted in PBS to the desired concentration for inoculation.

### Preparation of *Galleria* Moth Larva

*Galleria mellonella* larvae were acquired from Waxworms (waxworms.com). After delivery, larvae were portioned out into 100 mm Petri dishes, with 25-50 larvae per dish. Larvae weighing between 180–250 mg per larva and exhibiting ideal coloration (i.e. light tan with minimal or no spots or markings) were selected. Larvae were then stored at 37°C overnight to ensure they were starved for 24 hours prior to infection.

### *Galleria* Larva Infection

*Galleria* larvae were infected with a 10 µL inoculum of bacterial suspension, diluted to the designated OD600 as described above. Following dilution, 100 µL of the bacterial suspension was plated out onto LBA without antibiotics to determine the actual inoculum used. Plates were incubated at 37°C overnight, and colony-forming units (CFUs) were counted after 24–48 hours of incubation, depending on growth rate. Using the same inoculum, larvae were infected by injecting 10 µL into the left anterior proleg using a 27-gauge needle attached to a 1mL syringe. The injection was done using a KD Scientific syringe pump set to 7.20 mL/hour to deliver a total volume of 0.01 mL. Following inoculation, the larvae were moved to 37°C and monitored every 12 hours to assess mortality. Dead larvae were removed from the population. For antibiotic treatment, after a 4-hour incubation at 37°C, 10 µL of imipenem suspended in PBS was injected into the right anterior proleg of the larva. Larvae were then monitored at 8 hours post-treatment and every 12 hours thereafter.

### Measurement of bacterial CFU inside hemolymph

Following the *Galleria* infection, five *Galleria* larvae were collected from the live larvae every 24 hrs starting from 0 to 96 hours post-infection (hpi). The larvae were transferred individually to a - 20°C freezer for 3 minutes or until they became inactive (anesthetized). Larvae were then submerged in prechilled 70% ethanol for 20 seconds for sterilization and briefly dried on a sterile towel. The base of the larva was removed using sterile surgical scissors, and the hemolymph was gently squeezed out into a 1.5 mL tube containing 100 µL hemolymph anticoagulant solution (26 mM sodium citrate, 30 mM citric acid, 100 mM glucose, 140 mM NaCl, pH 4.11) (30) using blunt-ended forceps. The tube was spun down at 200 x g for 10 minutes, resuspended in 100 µL anticoagulant solution, and plated to determine the total CFU. To assess adherent and internalized bacteria per hemocyte, pelleted cells were resuspended in 200 µL anticoagulant and the wash step was repeated 2 times. The cell suspension was split into two aliquots: one aliquot contained no antibiotics to measure total cell-associated bacteria, i.e., adherent and internalized bacteria, and the other was treated with 50 μg/mL polymyxin B for 1 hour at room temperature to kill off bacteria outside hemolymph and quantify internalized bacteria. The number of hemocytes per mL was calculated using a hemocytometer and trypan blue staining. Each aliquot was then plated on LBA containing 20 μg/mL chloramphenicol.

### Tn7 Fluorescent labeling of *Achromobacter xylosoxidans*

Fluorescent labeling of *Ax* strains was conducted by Tn7 mutagenesis. The strain GN050 was first grown overnight at 37°C in 3mL LB without antibiotics. Concurrently, two strains of S17λpir containing either pGPTn7::GFP (NovoPro) or pUXBF13 (Addgene) were grown overnight in 3mL LB supplemented with 100 μg/mL carbenicillin. Overnight cultures were spun down at 6000 rpm for 5 minutes, and the supernatant was removed. Cell pellets were resuspended in 1 mL LB, mixed at a 1:1:1 ratio, and spun down at 6000 rpm for 5 minutes. The resulting pellet was resuspended in 100 µL LB, spot-plated onto LBA and incubated for 48 hours at 37°C. Single colonies of *Ax* were streaked onto LBA supplemented with kanamycin (20 μg/mL) and chloramphenicol (100 μg/mL). Insertion of *gfp* genewas confirmed by PCR using the screening primers OHL706 (GCAGGAAAGAAACGTCGCGGGT) and OHL707 (ATTTCACATCTTTCTTTCCG). The presence of GFP-producing *Ax* colonies was confirmed by visualizing the bacteria under a fluorescent microscope and GFP^+^ colony was chosen for downstream fluorescence microscopy.

### Fluorescence Microscopy of hemolymph

Prior to hemolymph collection, coverslips were sterilized in 70% ethanol and allowed to air dry. Sterilized coverslips were placed into a 6-well plate and treated with 1 mL of 0.01% poly-L-lysine for 10 minutes at room temperature. The coverslips were carefully removed using sharp-ended forceps and excess poly-L-lysine solution was removed. They were washed 3 times with 1 mL PBS and transferred to a covered container at 4°C overnight to dry. *Galleria* hemolymph was collected as noted above, except that the wash solution was supplemented with 1 µL/mL CellMask Deep Red and 0.0734 µL/mL Hoechst 33432. Following the initial spin at 200 x g, the pelleted cells were washed twice with 500 µL anticoagulant solution. The resulting cell suspension was applied to the prepared coverslips, spun onto the coverslip at 200 x g for 10 minutes and allowed to settle for 10 minutes at room temperature. Excess material was removed, and the coverslips were washed 3 times with 2 mL PBS. Cells were fixed with 1 mL 4% paraformaldehyde (PFA) for 10 minutes at room temperature, washed 3 times with PBS, and incubated with 1 mL 125 mM glycine for 10 minutes. Excess glycine was removed, and coverslips were mounted on slides using Prolong glass antifade mountant. Cells were imaged using a NIKON *AX*R fluorescent confocal microscope at 60X magnification using DAPI, TRITC, and Cy5 channels. Images were acquired as Z-stacks (0.2 µm slices, 1024 x 1024 pixels). Final images were generated using maximum-intensity Z-axis projections for each channel. Image processing was done using ImageJ. For all images, maximum and minimum brightness levels were adjusted to the same threshold to remove background prior to generating composite images. Cell outlines were collected by softening the cell membrane channel three times, followed by image thresholding, after which the resulting outline was obtained.

### Mouse infections

To assess mouse survival following *Achromobacter xylosoxidans* infection, we employed an intratracheal instillation model as previously described (13) using 8-week-old C57BL/6J male mice. Following infection, mice were monitored every 8 hours for weight loss, activity level, and physical appearance. Once mice returned to a clinical score of ≤ 2, the observation period was extended to every 12 hours. Mice with a total clinical score of ≥ 5 or a score of 3 in any single category were considered moribund and euthanized.

### Statistical Analysis

Statistical assessment of infection data was performed using Kaplan-Meier survival analysis with multiple-comparison correction done using the Benjamini-Hochberg test to control the false discovery rate. Statistical analyses of other data were performed using one-way ANOVA with Dunnett’s multiple comparisons test, unless otherwise noted. All statistical analyses were performed using GraphPad Prism software version 10.

## Results

### Survival of *Galleria mellonella* larvae infected with *Achromobacter xylosoxidans* recapitulates infection outcomes observed in mouse models

To investigate whether *Galleria mellonella* is an effective model for assessing disease severity of *Ax* infection, we compared *Galleria* larval survival with murine survival using the clinical isolates GN050 and GN008. GN050 and GN008 were previously characterized as cytotoxic and non-cytotoxic isolates, respectively (31), and have been used in murine infection models to investigate pathogenesis and immune responses during respiratory infection (12, 13). We therefore used these strains for the initial validation of *Ax* infection in *Galleria*.

We performed an intratracheal infection of C57BL/6J mice with *Ax*. 8-week-old mice were inoculated with either 5 x 10^7^ CFU, 1 x 10^7^ CFU, or 5 x 10^6^ CFU and monitored for up to 96 hours post infection (hpi). Mice infected with GN050 at 5 x 10^7^ CFU showed extremely severe symptoms, requiring euthanasia at the highest inoculum by 30 hpi (Figure 1A, Figure S1A-C, P). Infection with either 1 x 10^7^ CFU or 5 x 10^6^ CFU of GN050 resulted in ≥ 50% mortality by 60 hpi. GN008, by contrast, showed substantially less severe disease and required the highest dose 5 x 10^7^ CFU to significantly reduce mouse survival, with the majority of mice requiring euthanasia due to weight loss rather than elevated overall clinical scores (Figure 1B, Figure S1D-F, Q).

**Figure 1.**
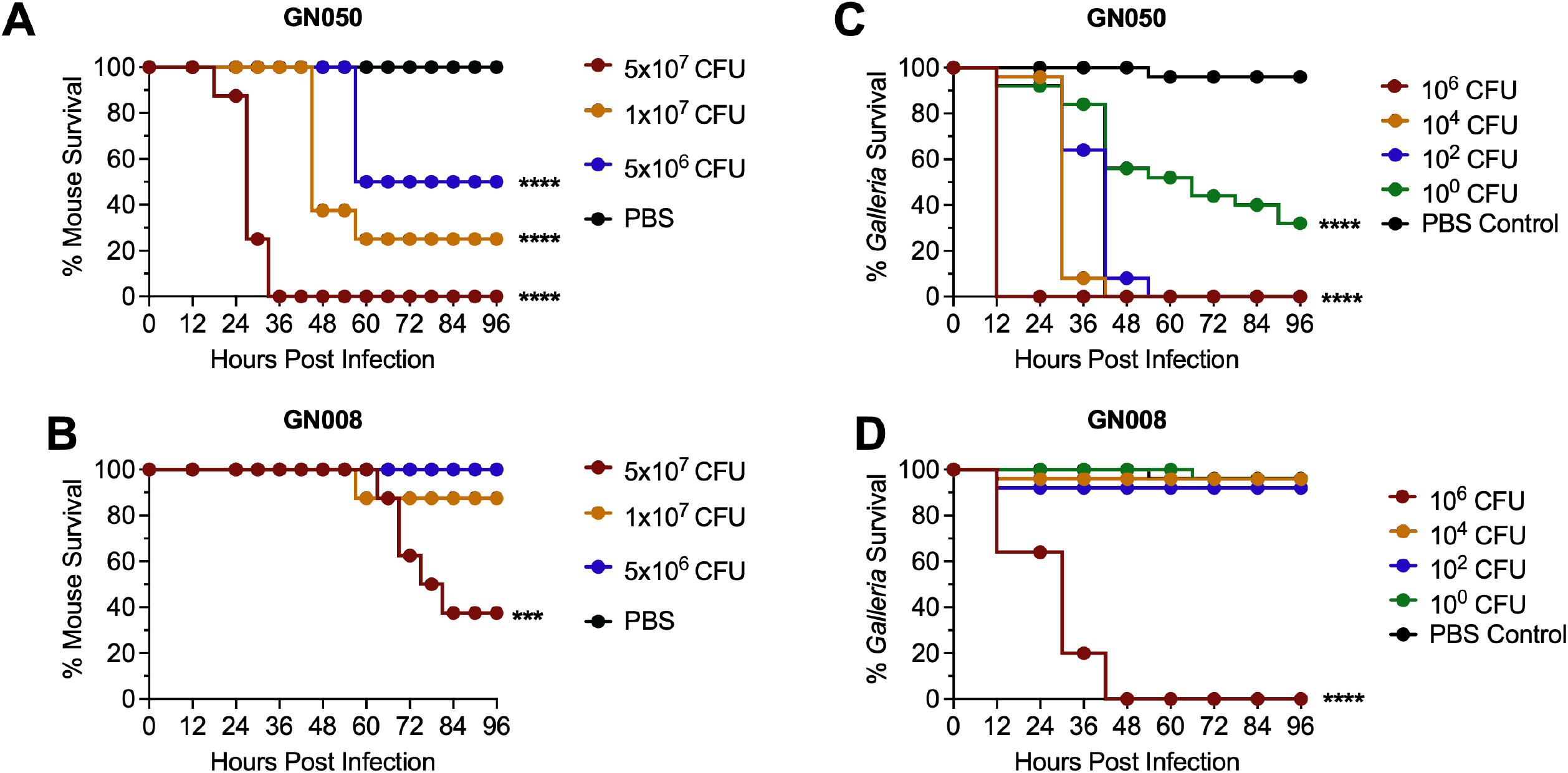
Survival of mice and *Galleria mellonella* larvae infected with *Achromobacter xylosoxidans* GN050 and GN008. Eight-week-old mice were infected with different doses of GN050 **(A)** and GN008 **(B)** by intratracheal instillation; n = 8. *Galleria mellonella* larvae were infected with different doses of GN050 **(C)** and GN008 **(D)**; n = 25. Survival was monitored up to 96 hours post infection (hpi). P-values were calculated using Kaplan-Meier survival analysis with correction for multiple comparisons by controlling the False Discovery Rate. ***: P<0.001, ****: P<0.0001.

We then infected *Galleria mellonella* larvae with GN050 and GN008. The procedure was outlined in Figure S2. *Galleria* larvae infected with GN050 required only approximately 1 CFU to achieve about 60% mortality by 96 hpi (Figure 1C). By contrast, GN008 did not result in substantial mortality until the highest dosage tested (1 x 10^6^ CFU) which resulted in 100% mortality by 48 hpi (Figure 1D). These results suggest that survival outcomes in the *Galleria* infection model followed the same trend as those observed in murine models.

### The *Galleria* infection model reveals a disconnect between *in vivo* virulence and *in vitro* cytotoxicity

We have access to NIH clinical isolates from previously published work (9, 29). Among them, the NIH-018 series was of particular interest. NIH-018-3, NIH-018-2, and NIH-018-1 were collected roughly one year apart, with NIH-018-3 collected first and NIH-018-1 collected last. *In vitro* characterization revealed a high degree of cytotoxicity among these isolates, which appeared to increase over time, and exceeded what was observed in GN050 (9). Thus, we were interested in further investigating their *in vivo* virulence. Following the same methodology as with GN050 and GN008, we infected *Galleria* larvae with the NIH-018 series clinical isolates (Figure 2A-C). Surprisingly, the most cytotoxic isolate, NIH-018-1, was by far the least virulent of the strains tested and showed similar mortality to GN008. The second biological replicate exhibited the same results. These findings suggest that *in vivo* virulence can diverge from *in vitro* cytotoxic phenotypes.

**Figure 2.**
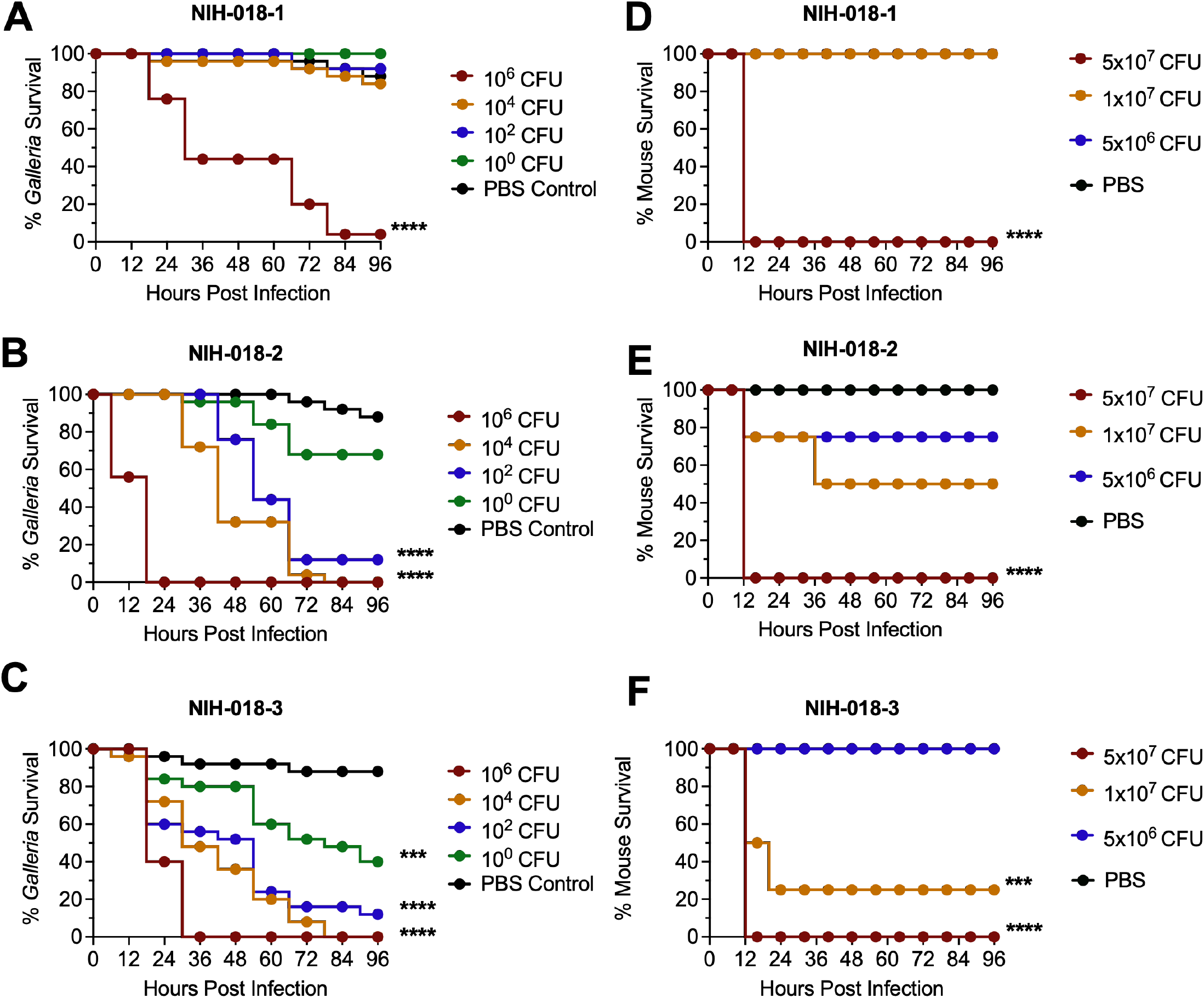
Survival of mice and *Galleria mellonella* larvae infected with *Achromobacter xylosoxidans* NIH-018 series. *Galleria mellonella* larvae were infected with different doses of NIH-018-1 **(A)**, NIH-018-2 **(B)**, and NIH-018-3 **(C)**; two biological replicates, n = 25/replicate. Eight-week-old mice were infected with different doses of NIH-018-1 **(D)**, NIH-018-2 **(E)**, and NIH-018-3 **(F)** by intratracheal instillation; n = 4. Survival was monitored up to 96 hpi. P-values were calculated using Kaplan-Meier survival analysis with correction for multiple comparisons by controlling the False Discovery Rate. ***: P<0.001, ****: P<0.0001.

To further validate whether the survival trend observed in *Galleria* reflected outcomes in murine models, we infected mice with the NIH-018 series. The results showed that infection of mice with the NIH-018 series exhibited survival patterns similar to those observed in *Galleria* (Figure 2D-F and Figure S1G-O, R-T). All mice infected with NIH-018 series isolates at the 5 x 10^7^ CFU inoculum showed extremely severe symptoms by 16 hpi, necessitating euthanasia as a result of morbidity (Figure S2G-O, R-T). NIH-018-1 infection resulted in no survival at 5 x 10^7^ CFU inoculum, but no morbidity resulting from 1 x 10^7^ CFU or 5 x 10^6^ CFU inoculum (Figure 2D). NIH-018-2 demonstrated an intermediate level of morbidity in mice with no survival at 5 x 10^7^ CFU inoculum, and no statistically significant reduction in survival at the 10^7^ CFU inoculum (Figure 3E). NIH-018-3, by contrast, showed the most severe infection with significant morbidity following infection at 1x10^7^ and 5 x 10^7^ CFU (Figure 3F). Assessment of the clinical scores and body weight indicated greater disease severity at the onset of infection in our NIH-018 strains compared to GN050 or GN008 (Figure S2). Overall, infection outcomes in mice infected with *Ax* isolates mirrored those observed in *Galleria*, while both differed markedly from trends observed in *in vitro* assays.

**Figure 3.**
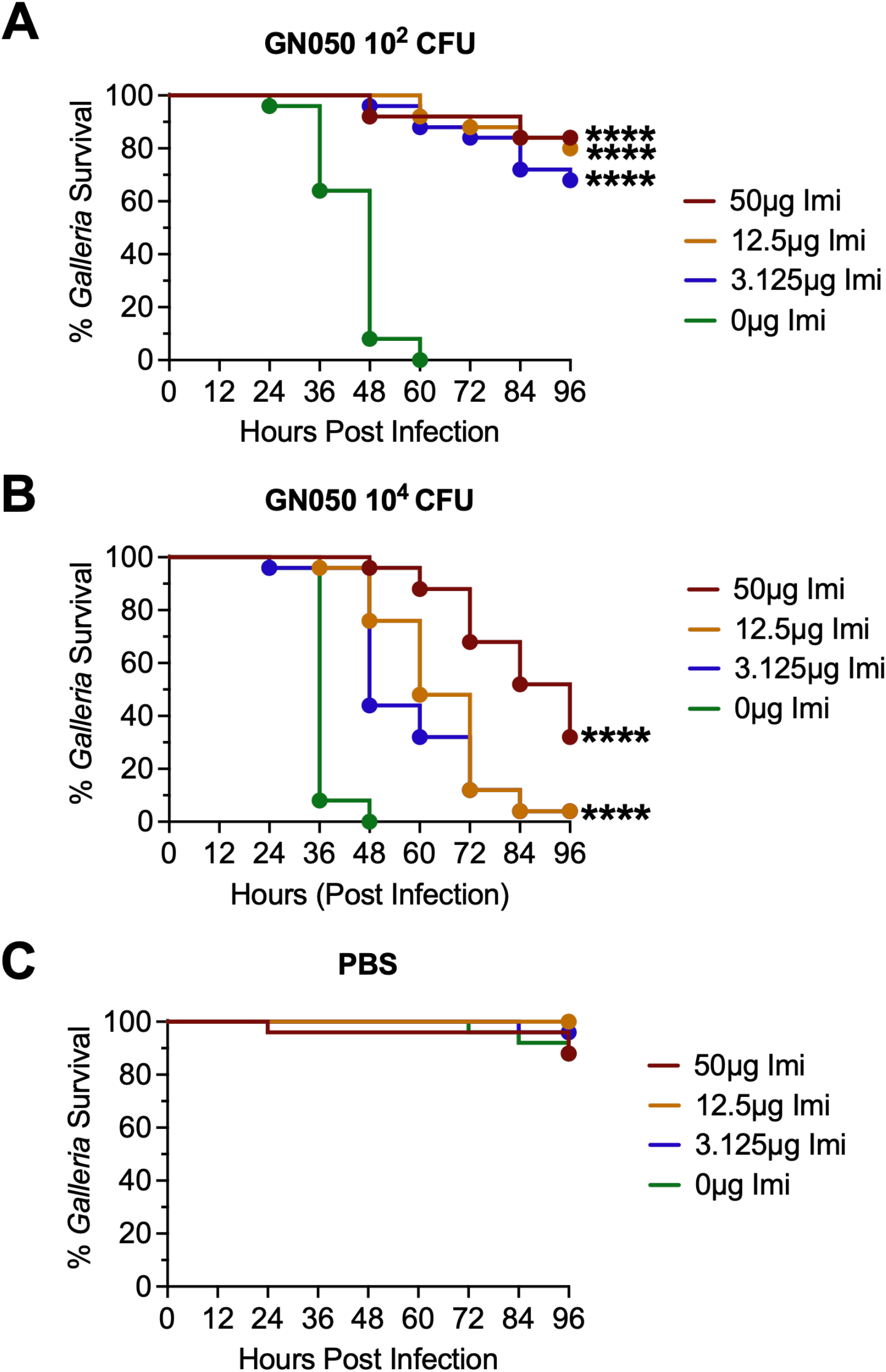
Survival of infected *Galleria mellonella* larvae under imipenem (IMI) treatment. *Galleria* larvae were infected with 10^2^ CFU **(A)** or 10^4^ CFU **(B)** of GN050, then treated with different doses of IMI at 4 hpi. Survival was monitored up to 96 hpi. **(C)** PBS control was injected to *Galleria* followed by injection of IMI at 4 hpi for toxicity assessment. P-values were calculated using Kaplan-Meier survival analysis with correction for multiple comparisons by controlling the False Discovery Rate. ****: P<0.0001.

### Antibiotic treatment reduces *Achromobacter*-induced mortality in *Galleria mellonella*

*Ax* is known to be highly resistant to many antibiotics, posing a significant challenge for patient treatment (9, 32) and highlighting the urgent need for novel antimicrobial discovery and development. *Galleria* has been employed as a model to evaluate antibiotic efficacy against other pathogens, such as *Acinetobacter* and *Pseudomonas* (23, 33-35). Antibiotics are known to exhibit different effects *in vivo* compared to *in vitro* (36). Thus, to facilitate the development of novel antimicrobial treatment strategies, we examined whether *Galleria* could be used to evaluate the *in vivo* efficacy of antibiotics against *Ax*.

*Galleria* larvae were infected with the strain GN050 at inocula of 1 x 10^2^ CFU and 1 x 10^4^ CFU. At 4 hpi, larvae were treated with imipenem (IMI), a broad-spectrum carbapenem used clinically to treat *Ax* infections (9). IMI-treated infected larvae showed a substantial reduction in mortality, with survival extending to at least 4 days post-infection at both GN050 inocula when treated with imipenem at 12.5 μg/mL or higher (Figure 3A, B). As a control, uninfected larvae treated with IMI showed no larval death, indicating that the antibiotic concentrations tested were not toxic to the larvae (Figure 3C). These results suggest that *Galleria* is a promising model for evaluating the *in vivo* effectiveness of antimicrobial agents against *Ax*.

### Growth of *Achromobacter xylosoxidans* and its association with hemocytes in *Galleria mellonella*

We monitored the total CFUs in larval hemolymph over time to evaluate how effectively *Galleria* larvae clear infection (Figure 4A, B). We observed an initial expansion of GN008 at 24 hpi, corresponding to a 10-fold increase and over a 1000-fold increase for GN050. Subsequent reduction in CFUs in GN008-infected larvae indicated clearance of bacteria from the hemolymph. GN050 showed a slower reduction with CFUs remaining 100-fold above the inoculum at 72 hpi.

**Figure 4.**
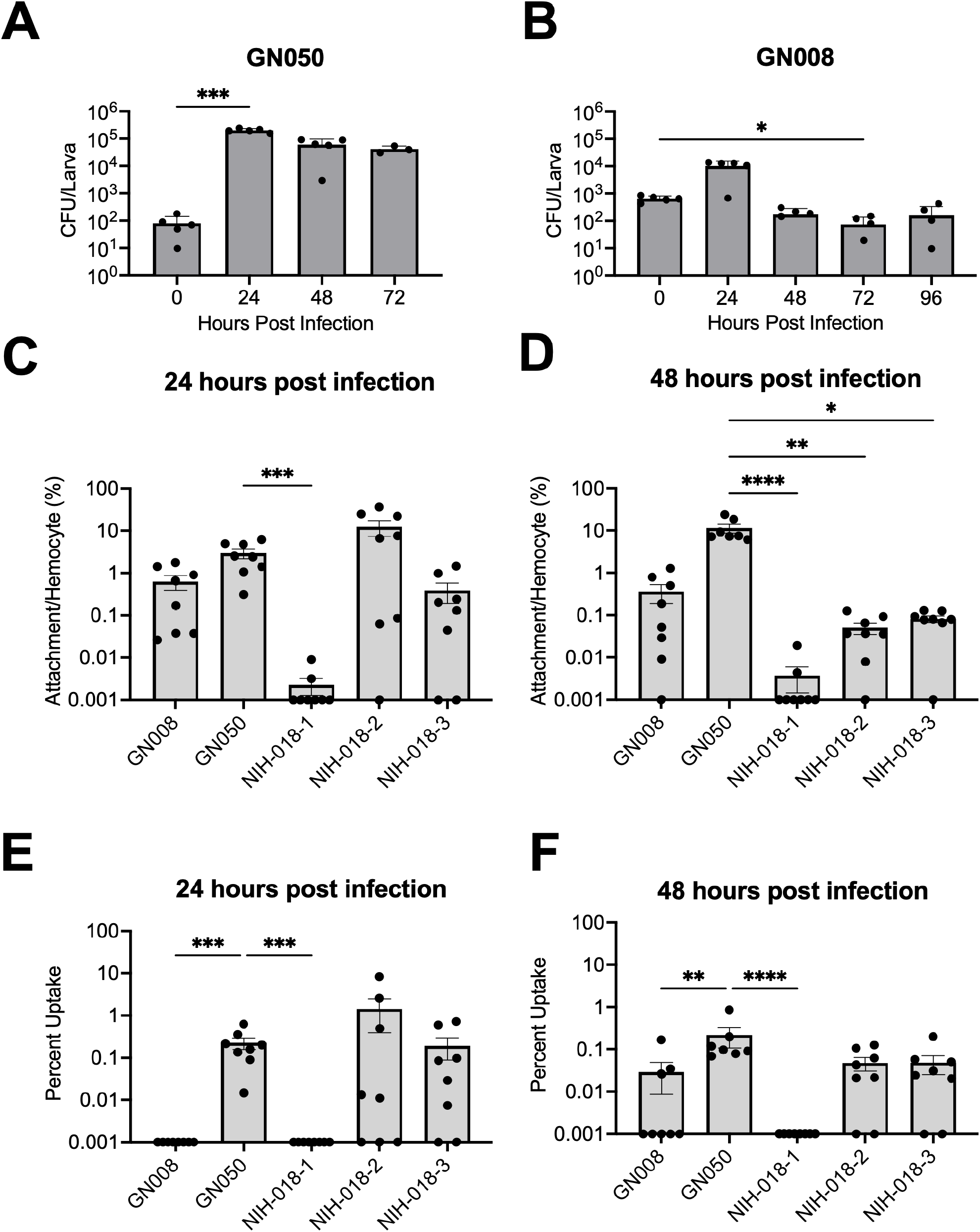
Growth and association of *Achromobacter xylosoxidans* in *Galleria. Galleria* larvae were infected with GN050 **(A)** and GN008 **(B)** at 10^2^ CFU and 10^3^ CFU, respectively. Bacterial growth (CFU) in hemolymph was monitored every 24 hours until 96 hpi; n = 5. All GN050 infected larvae were either dead or euthanized by 72 hpi. *Galleria* larvae were infected with different *Ax* isolates, including GN050, GN008, NIH-018-1, NIH-018-2 and NIH-018-3 at 10^3^ CFU. *Ax* attachment to hemocytes **(C-D)** and internalization into hemocytes **(E-F)** were monitored at 24 and 48 hpi; n = 8. P-values were determined by nonparametric one-way ANOVA with Dunn’s multiple comparisons test, comparing other time points to time 0 in (A, B) or comparing other isolates to GN050 in (C-F). *: P< 0.05, **: P< 0.01, ***: P<0.001, ****: P<0.0001.

Since *Ax* has been shown to adhere to and be internalized by macrophages *in vitro* (9, 12), we investigated whether *Ax* can similarly adhere to and become internalized by *Galleria* hemocytes. In addition, it was also shown that different *Ax* isolates can behave differently (9, 12). Thus, *Galleria* larvae were infected with different *Ax* isolates, GN050, NIH-018-2, and NIH-018-3 at 1 x 10^2^ CFU and GN008 and NIH-018-1 at 1 x 10^4^ CFU. These doses were chosen to ensure sufficient survival for monitoring of *Galleria* at 24 and 48 hpi, starting from a population of 30 larvae per condition (Figure 4C-F). Measurement of bacterial attachment to hemocytes revealed an overall low percentage of bacteria associated with hemocytes, with the NIH-018 series showing lower attachment than GN050 at 48 hpi (Figure 4C, D), but comparable levels of uptake, with the exception of NIH-018-1 (Figure 4E, F). NIH-018-1 exhibited neither attachment nor internalization. GN008 was also not internalized by hemocytes. These results suggest that internalization into hemocytes is associated with overall disease severity *in vivo*.

### Localization of *Ax* during *Galleria* infection

In order to verify the presence of *Ax* in *Galleria* hemocytes, fluorescent imaging was performed on hemocytes collected at 24 hpi from larvae infected with a Tn7-GFP-labeled *Achromobacter* GN050. Among the collected hemocytes, very few cells appeared to contain bacteria, consistent with our observations when measuring CFUs per hemocyte (Figure 4). Among the hemocytes observed to harbor bacteria, the majority appeared to be morphologically similar to oenocytoids (Figure 5, Cell Type 1), which are known to be phagocytic (37). Additional cell morphologies consistent with granulocyte-like (Figure 5, Cell Type 2) and plasmatocyte-like hemocytes (Figure 5, Cell Type 3) were also identified. Granulocytes, which function through lysis or degranulation to promote plasmatocyte binding, did not appear to contain *Ax* bacteria. In contrast, plasmatocytes were observed to contain bacteria, a finding consistent with previous studies (37, 38). These results suggest that *Ax* is internalized by phagocytic innate immune cells in *Galleria*, analogous to what occurs during vertebrate infection.

**Figure 5.**
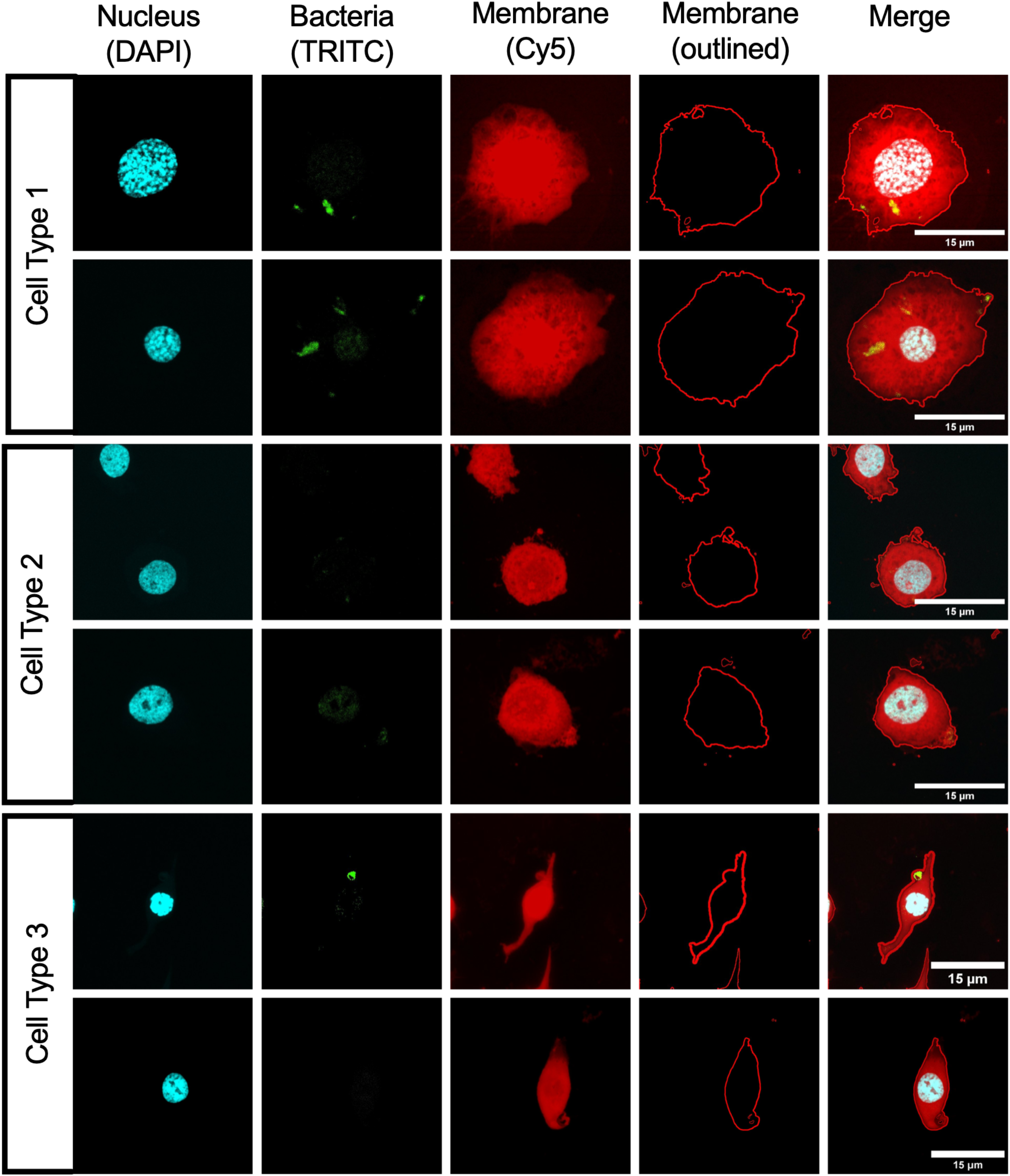
Uptake of *Ax* by hemocytes. *Galleria* larvae were infected with GN050-GFP at 10^2^ CFU. At 24 hpi, hemocytes were collected, stained and visualized using fluorescence microscopy. Representative images of identified hemocyte populations were divided into Cell type 1: oenocytoids-like cells, Cell Type 2: granulocytes-like cells, and Cell Type 3: plasmatocytes-like cells. DAPI: cell nucleus, TRITC: GN050-GFP, Cy5: CellMask Deep Red plasma membrane stain. Membrane outlines and merged images are shown.

## Discussion

Investigation of the pathogenesis and virulence of novel pathogens can be expensive and ethically demanding due to the use of mice or other vertebrate models of infection. Here, we present an alternative model for studying *Achromobacter xylosoxidans* infection and antimicrobial discovery. *Galleria* larvae are substantially less expensive and raise fewer ethical considerations than vertebrate models, making them an excellent alternative *in vivo* model. Our *Galleria* infection model recapitulated the same survival trends observed in murine models (Figure 1), yet these *in vivo* trends diverged from the *in vitro* cytotoxicity data (Figure 2; (9)). Previous studies have suggested the similar trend that different clinical isolates may have disparities in infection dynamics and severity that are not correlated with *in vitro* assays (9, 13). Therefore, using *Galleria* as an *in vivo* model in virulence studies can mitigate the substantial costs and ethical burden associated with mouse models.

*Achromobacter* sp. is known for highly antibiotic-resistant phenotypes (32, 39). *Galleria* larvae have been utilized as a model for assessing the effectiveness of antibiotics *in vivo* against infections caused by other pathogens (23, 35). Here, we validated that *Galleria* can serve as an effective *in vivo* model for evaluating antibiotic treatment against *Ax* (Figure 3). Beyond antibiotic testing, *Galleria* has also been utilized to characterize the *in vivo* effectiveness of virulence inhibitors (22, 40). The utility of such a model becomes apparent in the context of *Ax* virulence factors, such as ArtA, an RTX adhesin important for adherence and cell invasion (13). The design of small molecules to inhibit adhesins and other virulence factors to reduce disease severity may be a promising area of research, in which the *Galleria* model can be used to assess *in vivo* efficacy. In short, the *Galleria* model can serve as an initial *in vivo* model to test the effectiveness of antimicrobials, facilitating novel drug discovery and development.

Cytotoxic *Ax* isolates are known to kill innate immune cells such as macrophages, while non-cytotoxic isolates are less capable of causing cell death (9, 28). In the *Galleria* infection model, GN050 (a cytotoxic isolate) was able to multiply rapidly by 24 hpi and maintain a high number of bacteria inside Galleria at later time points, although the CFU counts dropped after 24 hpi. The reduction in GN050 growth could be due either to immune-mediated bacterial killing or to nutrient limitation resulting from the high bacterial density inside *Galleria*. Given the ∼1000-fold increase in GN050 bacterial numbers, the latter explanation may be more likely. In contrast, GN008 showed limited replication and a significant reduction in bacterial counts inside *Galleria* by 72 hpi, suggesting rapid and effective clearance of GN008 by *Galleria*. Since *Galleria* lacks an adaptive immune system, these findings suggest that innate immunity plays an important role in controlling *Ax* infection; however, more cytotoxic and virulent isolates may be able to evade or counteract the innate immune system, leading to persistence or proliferation, indicating that adaptive immunity may contribute to host defense against *Ax* to avoid chronic infection.

Assessment of *Ax* adhesion to and internalization by *Galleria* hemocytes revealed an overall low level of bacterial association. GN050 attached to approximately 10% of hemocytes, while other isolates attached to less than 1% of the cells at 48 hpi (Figure 4). Uptake of GN050, NIH-018-2, and NIH-018-3 was comparable and occurred in around 0.1% of hemocytes while uptake of GN008 and NIH-018-1 was significantly lower than that of the other isolates. In agreement with bacterial uptake trends, disease severity in mice and *Galleria* was much higher for GN050, NIH-018-2 and NIH-018-3 compared to the other two isolates. These observations suggest that uptake of *Ax* by phagocytic cells is associated with disease severity. Moreover, in our previous *in vitro* studies (9), attachment and uptake among NIH-018 isolates were similar; however, they differed in the *Galleria* infection model, with NIH-018-1 exhibiting the least uptake. Moreover, NIH-018-1 is the most cytotoxic isolate *in vitro*, but causes the least severe infection among the NIH-018 isolates. These results highlight the complexity of host-pathogen interactions *in vivo*, which could be missed in *in vitro* experiments.

To visualize internalized bacteria in hemocytes, we performed fluorescent imaging of GN050-GFP infecting *Galleria* larvae. Hemocytes were collected at 24 hpi and visualized. Oenocytoids, granulocytes, and plasmatocytes are the main phagocytic cells in *Galleria*, and granulocytes can take up *E. coli* at the fastest rate (37). From our *in vivo* samples, *Ax* was found inside oenocytoids and plasmatocytes, but not granulocytes (Figure 5). This discrepancy may reflect enhanced susceptibility of granulocytes to GN050-mediated killing or a pathogen-specific immune response. Further studies are needed to investigate which specific immune cells are critical for responding to *Ax*, which is beyond the scope of this study.

While our results strongly support the use of *Galleria* as an *in vivo* infection model, this study has limitations. Our work utilized a small number of mice for the exploratory infections with the NIH-018 series. However, this limitation itself underlines the value of *Galleria mellonella* as a preliminary *in vivo* model, given the cost and ethical implications of mouse work.

In summary, we have demonstrated that the *Galleria* infection model enables researchers to obtain important insights into disease severity and interactions between bacteria and the innate immune system. It can be a useful tool for the identification of novel virulence factors in *Ax* clinical isolates. In addition, the *Galleria* model may serve as an effective system for the preliminary screening and testing of novel antimicrobials.

## Figure legends

Supplemental Figure 1: Clinical scores and weight changes of mice infected with *Ax* over time. Eight-week-old mice were infected with GN050 **(A-C, P)**, GN008 **(D-F, Q)**, NIH-018-1 **(G-I, R)**, NIH-018-2 **(J-L, S)** and NIH-018-3 **(M-O, T)** and clinical scores and weight changes were monitored up to 96 hpi. Clinical scores were determined by summing scores from the following rubric: Physical appearance: 0=normal, Bright Active and Alert (BAR), 1=BAR, mild hunch and ruffled fur, abnormal stance, mild ocular or nasal discharge, 2=Quiet Alert and Responsive (QAR), hunched, squinted eyes, ears pinned, heavy ocular or nasal discharge, increased breathing. 3=Unresponsive, recumbency, eyes closed, severely hunched, labored breathing. Activity: 0=BAR, 1=BAR, abnormal gait, mildly lethargic, 2=QAR, reluctant to move, lethargic, 3=moribund/inability to move. Weight change: 0=<10%, 1=10-15%, 2=16-20%, 3=>20%. Any mouse with a clinical score of 5 or greater in total or 3 in any one category was categorized as moribund and euthanized.

Supplemental Figure 2: Schematic of *Galleria mellonella* infection. **(A)** Larvae were infected with 10μl of diluted bacteria and monitored every 12 hours, or **(B)** when treating *Galleria* with antibiotics, imipenem was administered at 4 hpi, and dead larvae were counted every 12 hours.

## Notes

### Competing Interest Statement

The authors have declared no competing interest.

