## Supplementary figures and images for "Establishment of *Galleria mellonella* as a Model for *Achromobacter xylosoxidans* Infection"

### Figure S1

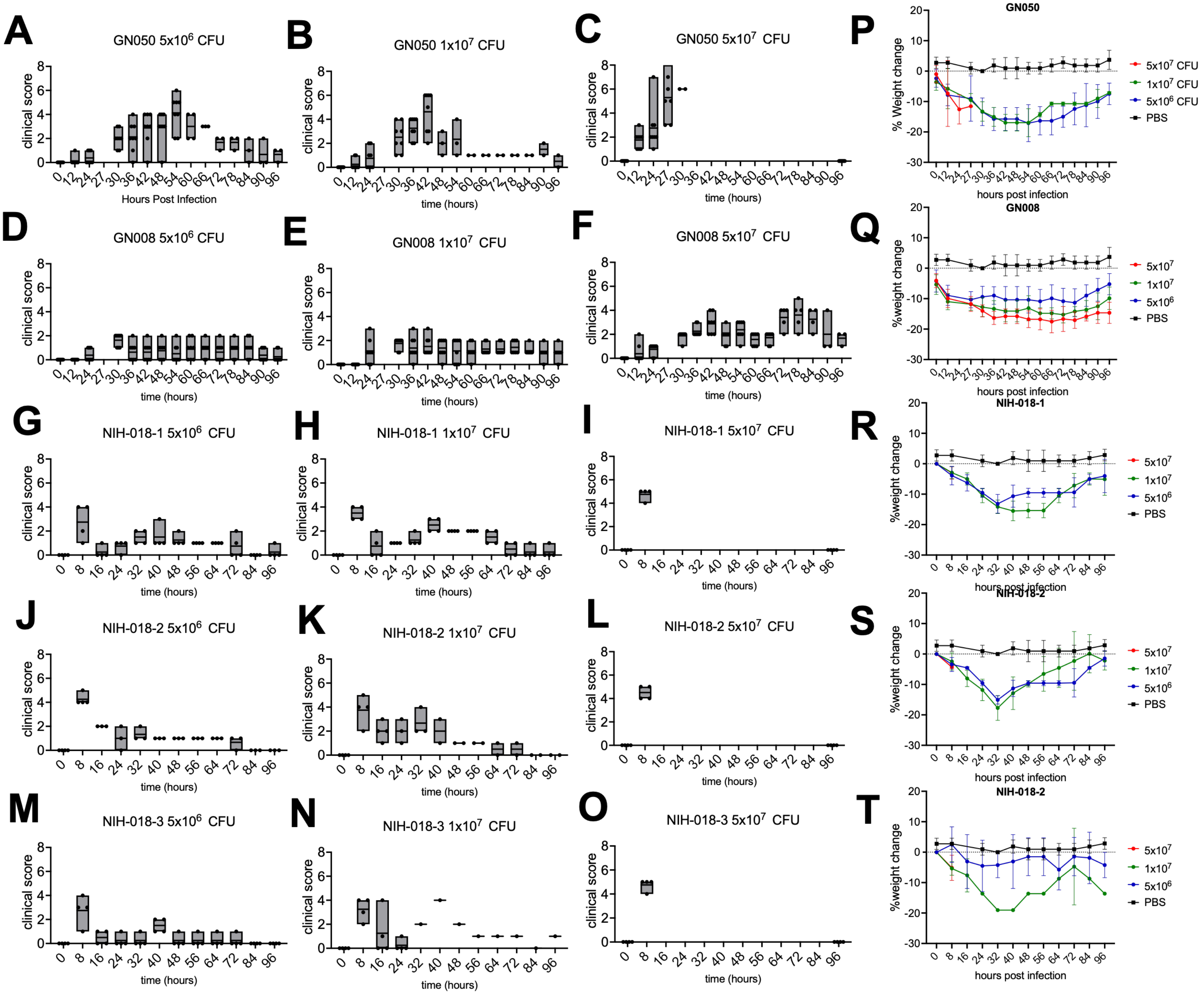

### Figure S2

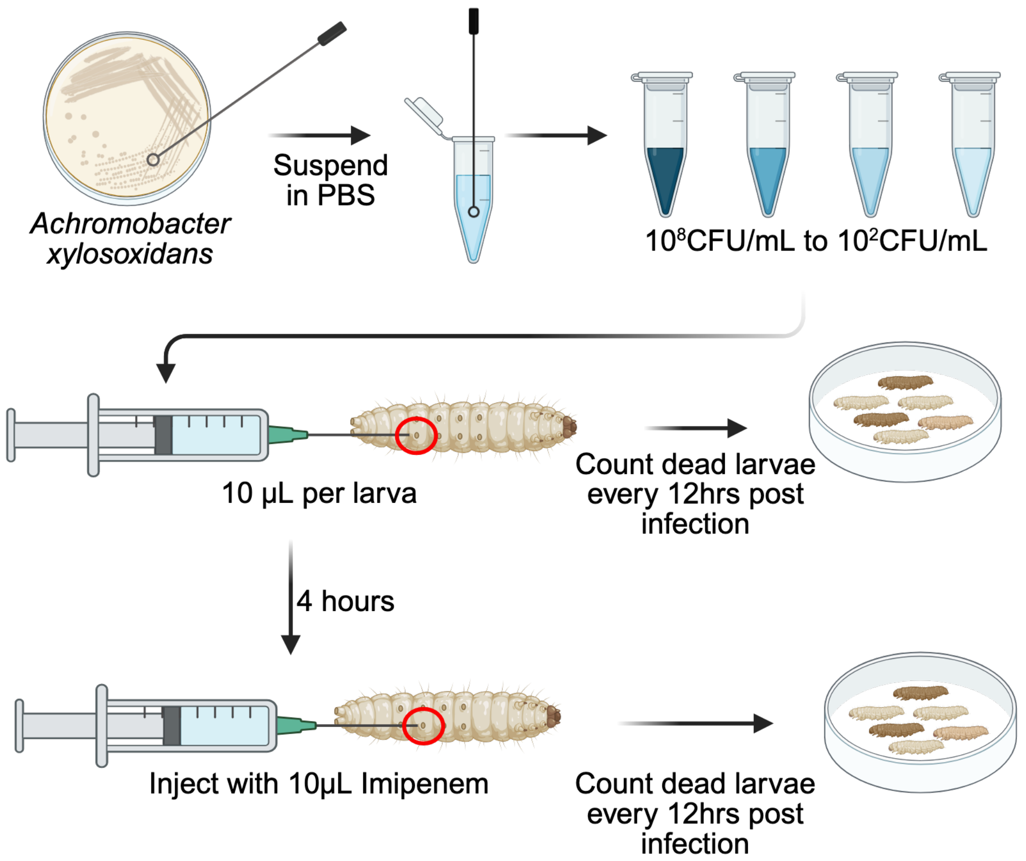
